# Logistic Regression Analysis of Multiple Interosseous Hand-Muscle Activities using Surface Electromyography during Finger-Oriented Tasks

**DOI:** 10.1101/278051

**Authors:** Masayuki Yokoyama, Masao Yanagisawa

## Abstract

Intrinsic hand muscles are densely located in the hand, and the myoelectric observation from the surface is sometimes unreliable because of some outside influences that may interfere with the signals. In the present study, we evaluated the activities of multiple interosseous hand-muscles which densely located in the hand, through analyzing the surface electromyographic signals during finger-oriented tasks using univariate and multivariate logistic regression models. Ten healthy subjects participated in our experiment, and isometrically exercised each finger one by one in flexed form. The result of a univariate analysis with the power and amplitude domain predictor variables of the surface electromyographic signals showed significant consistency between the activated finger and the inserted finger of the dorsal interosseous muscles to the proximal phalanx (*P* < 0.001). Meanwhile, the results of a multivariate analysis showed a higher correlation of the regression model of the fourth dorsal interosseous muscle during the action of the ring finger using frequency-domain variables (the Nagelkerke *R*^2^ = 0.716 when the median frequency was used), compared to the model without the frequency-domain variables (the Nagelkerke *R*^2^ = 0.583). Our result showed that the logistic regression models have a particular possibility for the analysis of the surface electromyographic signals of densely located hand-muscle activities related to the finger-oriented tasks.

## 1. Introduction

Surface electromyography (sEMG) is widely used to observe the state of muscle contraction in a non-invasive manner (Merletti and Parker, 2004). The characteristics of the sensor, which include safe use and less cost (Yokoyama et al., 2017), help analyze human actions not only for clinical applications, such as hand rehabilitation systems (Leonardis et al., 2015; Ho et al., 2011; Ang et al., 2009), but also for non-clinical applications in information technology development, such as hand-oriented user interface (Saponas et al., 2008; Zhang et al., 2009). Hand and finger gesture recognition is a well-studied field as a key technology of the hand-oriented user interface. Compared to the significantly heavier mechanical gloves (Popescu et al., 2000; Jack et al., 2001; Dovat et al., 2008; Ma et al., 2016; Demain et al., 2013), the light-weighted sEMG sensors potentially save the user with long-term use from exhaustion (Ma et al., 2016; Demain et al., 2013). Therefore, the development of light-weighted hand wearable devices using sEMG sensors is encouraged and fostered to create promising products.

Muscles related to hand and finger activities are located in the forearm (i.e., extrinsic hand muscles) and hand (i.e., intrinsic hand muscles) (Prerotto, 2011). sEMG electrodes are generally put on the forearm to analyze activities of the wrist (Szeto and Lin, 2011) and the whole hand, such as grasping motion (Green et al., 2015; Hoozemans and van Dieën, 2005). In case of finger-related activities, intrinsic hand muscles work harmoniously with extrinsic hand muscles (Johnston et al., 2010; Buford et al., 2005; Al-Sukaini et al., 2016; Lee and Wisser, 2012).

Johnston et al. (2010) reported the co-activation of EMG signals of extrinsic and intrinsic hand muscles during two-digit grasping motions using the thumb and the index finger with various wrist angles. Al-Sukaini et al. (2016) examined the dominance of extrinsic or intrinsic hand muscles in finger flexion using video motion tracking, and their results showed that 69.5% of the hands of the subjects had extrinsic dominance.

However, the extrinsic hand muscles themselves are not suitable to analyze the independent motion of each finger because the flexor digitorum superficialis muscle, as a flexor, and the extensor digitorum muscle, as an extensor, manipulate multiple fingers (i.e., from the index to the little fingers). The boundaries of muscle fibers for each finger are not explicitly separated, except for some pollicis, indicis, and digiti minimi muscles. On the other hand, intrinsic hand muscles are located very close to the fingers and each muscle is inserted into a single phalanx of a finger.

It is not necessary to put the electrodes on all related muscles to analyze a motion, because according to the phenomenon “muscle synergies” (d’Avella et al., 2003), a behavior produced by contraction of many muscles can be represented by a small subset. This is important especially in the field of a hand-oriented user interface, where a small and light-weighted design is required for a wearable device. Intrinsic-domain sensing has a potential to help the designers because the electrodes may only be put on the hand and the design around the forearm may not be considered. Research to utilize this phenomenon for the analysis of finger activities was conducted by Choi et al. (2010), who evaluated the finger force during pinching by the thumb and the index finger only from three intrinsic hand muscles.

However, there is a problem in measuring sEMG signals from intrinsic hand muscles because the area in contact with the electrodes is quite small, so the signals from the adjacent muscles can interfere. Therefore, relatively few studies of finger activities by sensing intrinsic hand muscles have been conducted so far (Choi et al., 2010). Choi et al. (2010) applied an artificial neural network (ANN) to predict the finger force, but the ANN classifiers are known as a black box (Ayer et al., 2010), which finds suitable features automatically without knowing the characteristics of the input data. Due to an overfitting problem, the results predicted by ANN seemed good. However, it was impossible to obtain clues to select an independent variable from the results, so the network had to be designed through trial-and-error.

In this study, we examined an availability of logistic regression analysis to evaluate activities of fingers by sensing sEMG signals obtained from densely located hand-muscles. We evaluated an isometric flexion form of each finger because the subject can easily control a single finger independently. We also examined which independent variables of the regression models would contribute to the analysis of the finger activities. Kaplanis et al. (2009) evaluated 13 independent variables extracted in the time, frequency, and bispectrum domains from the Biceps Brachii muscle generated under isometric voluntary contraction. They analyzed the data univariately using the Wilcoxon signed rank test. Finger activities, to which many extrinsic and intrinsic muscles are related, are supposed to be more complex than upper-limb activities, so we adopted logistic regression models for the multivariate analysis of finger activities. We focused on the binary outcome, which represents the state of finger activities as active or non-active, to simplify the analysis. We chose four dorsal interosseous muscles (DI) in a hand as the measurement targets of our experiment by putting the electrodes to measure the sEMG signals of the DI muscles on the back of the hand, which did not prevent any flexion and extension activities of the fingers in practical life.

## 2. Materials and Methods

### 2.1. Subjects and Experimental Environment

Ten healthy male subjects in their early twenties participated in our experiment. This research was non-clinical, and the observed data were analyzed anonymously; therefore, verbal consent was obtained from all participants. The study was approved by the Human Ethics Committee of Waseda University (No.2016-HN035). Our data is available at https://doi.org/10.6084/m9.figshare.6467801.

We used wireless sEMG sensors OE-WES1224 (Oisaka Electronic Equipment Ltd.) with bipolar Ag/AgCl electrodes. The inter-electrode distance was approximately 20 mm, and the diameter of the electrodes was 10 mm. A disposal sheet was cut into 10-mm squares to position the electrodes on the limited space of the back of the hand. The sampling frequency of the sensors was 1 kHz. A 16-bit A/D converter and a 4th-order band-pass filter from 30 to 700 Hz were used.

### 2.2. Experimental Procedure

The subjects were told to sit down on a chair and to hold their palms on a desk, as shown in Fig. 1. The angle of the elbow was almost perpendicular with a natural and relaxed posture. The subjects forced the desk with a single finger in a flexion form, and then relaxed with the same posture (i.e. isometric muscle contraction). The forcing and relaxing conditions were maintained for 3 s each, and iterated for 50 s for each finger. The subjects were told to force the desk with as much strength as possible (i.e. maximum voluntary contraction). The tips of the fingers were always in contact with the desk, and the motion was quasi-static. There were two flexion forms of the finger: opened and closed forms (Long and Brown, 1964), shown in Fig. 2. This time we adopted the opened flexion form because the results had little difference between the two forms in our previous research (Yokoyama et al., 2015). We adopted the electrode positions used in (Prerotto, 2011), at the back of the hand between the metacarpal bones.

**Figure 1.**
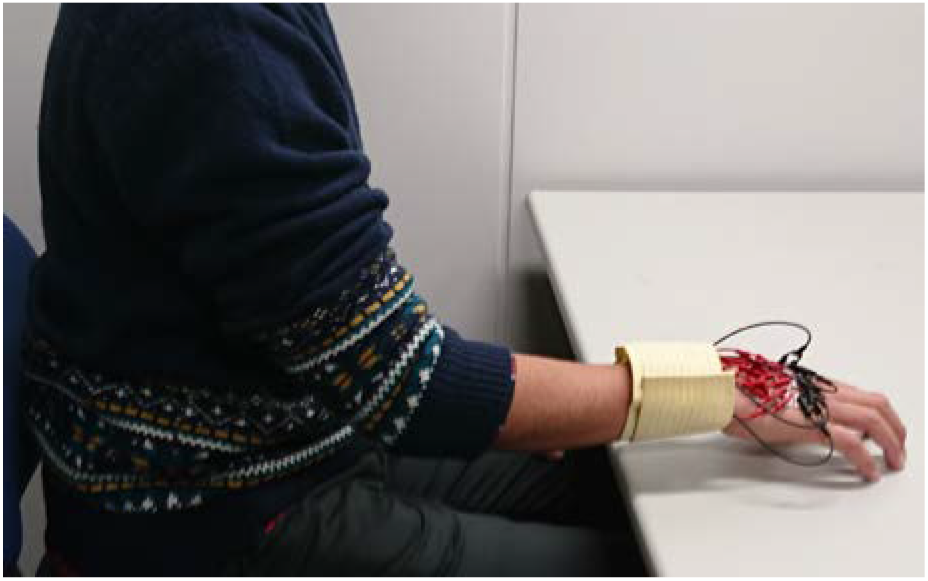
Pose of upper-limb of a subject in our experiment.

**Figure 2.**
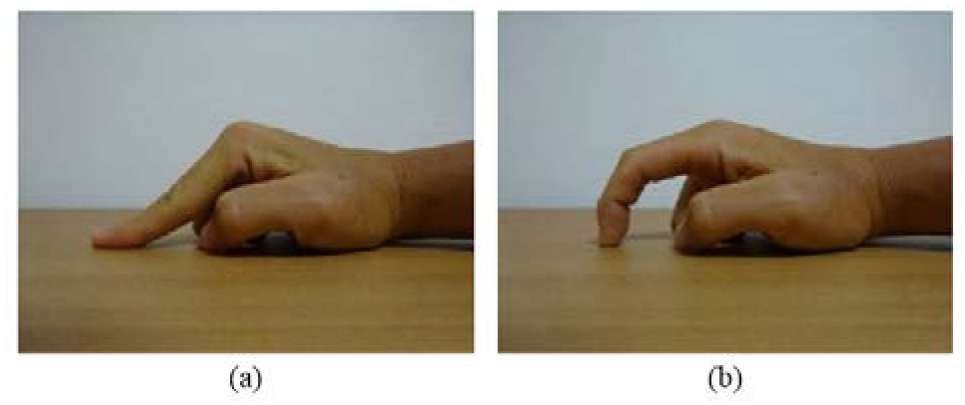
Two different forms of index-finger-flexion. (a) Opened form. (b) Closed form. (Yokoyama et al., 2015)

### 2.3. Data Analysis

Mean with standard deviation (SD), 95% confidence interval (CI), and intra-class correlation (ICC) of active (i.e., forced) and non-active (i.e., relaxed) data were calculated. Statistical analyses were performed using SPSS Statistics version 24.0 and Matlab for Windows.

We applied both univariate and multivariate logistic regression analyses to the data. The logistic regression model with multiple predictor variables was defined by the following equation:

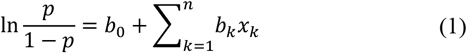

where *p* represents an outcome (or dependent variable) as a probability, *b*_0_…*b*_n_ represent the constant and the coefficients of the regression model, and *x*_1_…*x*_n_ represent the predictor (or independent) variables.

We chose the following six predictor variables: maximum amplitude (*V*_max_ [V]), mean amplitude (*V*_mean_ [V]), median frequency (*f*_med_ [Hz]), peak frequency (*f*_peak_ [Hz]), maximum power of the spectrum (*P*_max_ [V/Hz]), and total integrated power of the spectrum (*P*_total_ [V/Hz]).

*P*-value < 0.001 was considered to be statistically significant in our logistic regression analysis. We tested our regression models using mean values in the same experimental session of each subject, otherwise the variance of each predictor variable would be computed as much lower values.

## 3. Results and Discussion

### 3.1. Univariate Analysis

Table 1 shows the results of our univariate logistic regression analysis. The results of each predictor variable are shown as columns, with the correlated muscles to the finger activity shown as rows. The results during the forced state are shown as “active,” while those during the relaxed state are shown as “non-active.” Note that only the connected muscles to each finger are shown in Table 1, which include the first DI muscle inserted into the radial side of the index finger, the second DI muscle inserted into the radial side of the middle finger, the third DI muscle inserted into the ulnar side of the middle finger, and the fourth DI muscle inserted into the ulnar side of the ring finger, as shown in Fig. 3. The ICC of the active state was computed from the first seven datasets of each session of the related finger activity among all subjects. Meanwhile, the ICC of the non-active state of each muscle was computed from the first seven datasets of all sessions of the related and non-related finger activities among all subjects. The average values of signal-to-noise ratio (SNR) of each predictor variable are also shown in Table 1. The SNR of each subject as the linear scale was computed using the following equation:

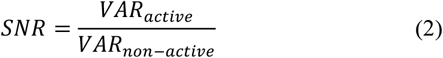
where *VAR_active_* represents the value of a predictor variable during the active state, and *VAR_non-active_* represents the value of a predictor variable during the non-active state.

**Figure 3.**
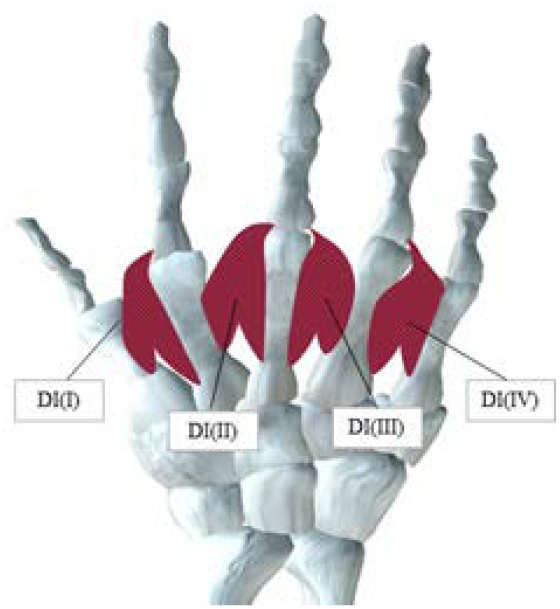
Interosseous muscles of the dorsal side of the right hand.

**Table 1.** Results of the univariate logistic regression analysis.

| Finger<br>Electrode<br>Position | Variable |  |  |  |  |  |
| --- | --- | --- | --- | --- | --- | --- |
| | $V_{\max}$ | $V_{\text{mean}}$ | $f_{\text{med}}$ | $f_{\text{peak}}$ | $P_{\max}$ | $P_{\text{total}}$ |
| Index |  |  |  |  |  |  |
| DI(I) |  |  |  |  |  |  |
| Non-Active |  |  |  |  |  |  |
| mean (SD) | $0.037 \pm 0.016$ | $0.007 \pm 0.002$ | $181.838 \pm 28.835$ | $93.963 \pm 48.003$ | $1.265 \pm 0.201$ | $0.139 \pm 0.054$ |
| CI | (0.0261 - 0.048) | (0.005 - 0.008) | (161.211 - 202.466) | (59.624 - 128.302) | (1.122 - 1.409) | (0.101 - 0.178) |
| ICC | 0.904 | 0.927 | 0.977 | 0.963 | 0.355 | 0.918 |
| Active |  |  |  |  |  |  |
| mean (SD) | $0.228 \pm 0.111$ | $0.041 \pm 0.020$ | $205.225 \pm 34.333$ | $164.636 \pm 53.617$ | $8.142 \pm 3.891$ | $1.072 \pm 0.562$ |
| CI | (0.149 - 0.308) | (0.027 - 0.056) | (180.664 - 229.785) | (126.281 - 202.992) | (5.359 - 10.926) | (0.670 - 1.474) |
| ICC | 0.982 | 0.995 | 0.993 | 0.918 | 0.984 | 0.995 |
| SNR | $5.423 \pm 6.119$ | $5.556 \pm 5.040$ | $1.201 \pm 0.152$ | $1.926 \pm 0.930$ | $5.134 \pm 4.229$ | $6.998 \pm 7.744$ |
| P | <0.001 | <0.001 | 0.054 | 0.003 | <0.001 | <0.001 |
| Middle |  |  |  |  |  |  |
| DI(II) |  |  |  |  |  |  |
| Non-Active |  |  |  |  |  |  |
| mean (SD) | $0.043 \pm 0.017$ | $0.007 \pm 0.002$ | $199.839 \pm 24.102$ | $122.574 \pm 56.240$ | $1.230 \pm 0.403$ | $0.162 \pm 0.067$ |
| CI | (0.031 - 0.055) | (0.005 - 0.008) | (182.598 - 217.080) | (82.342 - 162.806) | (0.941 - 1.518) | (0.114 - 0.210) |
| ICC | 0.919 | 0.926 | 0.977 | 0.954 | 0.876 | 0.931 |
| Active |  |  |  |  |  |  |
| mean (SD) | $0.202 \pm 0.105$ | $0.037 \pm 0.015$ | $233.447 \pm 23.053$ | $202.405 \pm 28.271$ | $7.015 \pm 2.931$ | $1.006 \pm 0.414$ |
| CI | (0.127 - 0.277) | (0.026 - 0.048) | (216.956 - 249.939) | (182.181 - 222.629) | (4.918 - 9.112) | (0.710 - 1.302) |
| ICC | 0.972 | 0.984 | 0.984 | 0.740 | 0.980 | 0.985 |
| SNR | $3.060 \pm 1.204$ | $3.949 \pm 1.551$ | $1.096 \pm 0.068$ | $1.603 \pm 1.411$ | $4.282 \pm 1.857$ | $4.468 \pm 2.099$ |
| P | <0.001 | <0.001 | 0.001 | 0.001 | <0.001 | <0.001 |
| DI(III) |  |  |  |  |  |  |
| Non-Active |  |  |  |  |  |  |
| mean (SD) | $0.026 \pm 0.013$ | $0.005 \pm 0.002$ | $195.459 \pm 14.484$ | $100.025 \pm 48.403$ | $0.852 \pm 0.268$ | $0.104 \pm 0.046$ |
| CI | (0.017 - 0.035) | (0.004 - 0.006) | (185.098 - 205.820) | (65.400 - 134.651) | (0.660 - 1.044) | (0.071 - 0.138) |
| ICC | 0.944 | 0.923 | 0.927 | 0.936 | 0.668 | 0.949 |
| Active |  |  |  |  |  |  |
| mean (SD) | $0.215 \pm 0.131$ | $0.040 \pm 0.025$ | $225.525 \pm 18.692$ | $183.459 \pm 29.035$ | $7.558 \pm 5.008$ | $1.078 \pm 0.663$ |
| CI | (0.121 - 0.309) | (0.022 - 0.058) | (212.154 - 238.896) | (162.689 - 204.230) | (3.975 - 11.140) | (0.604 - 1.552) |
| ICC | 0.979 | 0.990 | 0.972 | 0.509 | 0.984 | 0.990 |
| SNR | $7.053 \pm 4.126$ | $7.321 \pm 4.089$ | $1.057 \pm 0.103$ | $1.615 \pm 0.993$ | $8.758 \pm 4.858$ | $8.965 \pm 5.088$ |
| P | <0.001 | <0.001 | 0.001 | <0.001 | <0.001 | <0.001 |
| Ring |  |  |  |  |  |  |
| DI(IV) |  |  |  |  |  |  |
| Non-Active |  |  |  |  |  |  |
| mean (SD) | $0.029 \pm 0.009$ | $0.006 \pm 0.002$ | $175.130 \pm 24.321$ | $81.499 \pm 44.176$ | $1.041 \pm 0.321$ | $0.116 \pm 0.041$ |
| CI | (0.022 - 0.035) | (0.005 - 0.007) | (157.732 - 192.529) | (49.897 - 113.100) | (0.811 - 1.270) | (0.086 - 0.145) |
| ICC | 0.901 | 0.960 | 0.970 | 0.962 | 0.904 | 0.943 |
| Active |  |  |  |  |  |  |
| mean (SD) | $0.260 \pm 0.183$ | $0.047 \pm 0.033$ | $213.428 \pm 20.391$ | $173.584 \pm 40.373$ | $9.394 \pm 7.444$ | $1.234 \pm 0.811$ |
| CI | (0.129 - 0.391) | (0.024 - 0.070) | (198.841 - 228.014) | (144.703 - 202.465) | (4.069 - 14.719) | (0.654 - 1.814) |
| ICC | 0.979 | 0.990 | 0.977 | 0.748 | 0.983 | 0.988 |
| SNR | $7.817 \pm 4.77$ | $7.900 \pm 5.906$ | $1.244 \pm 0.265$ | $2.263 \pm 0.934$ | $8.821 \pm 6.527$ | $10.805 \pm 8.052$ |
| P | <0.001 | <0.001 | 0.001 | <0.001 | <0.001 | <0.001 |

The active and non-active states were significantly discriminated using the amplitude (*V*_max_ and *V*_mean_) and power (*P*_max_ and *P*_total_) of the sEMG signals, hence acting as the predictor variables. However, the two states were not significantly discriminated using the frequency-domain variables (*f*_med_ and *f*_peak_). In fact, none of the *f*_med_ values was recorded as significant in our experiment, but only the two states of the third and fourth DI muscles were discriminated using *f*_peak_, making it a potential predictor variable for our univariate logistic regression analysis as well.

Some of the ICCs using *f*_peak_ and *P*_max_ as the predictor variables showed fairly low values. The ICC of the third DI muscle using *f*_peak_ was 0.509 when the middle finger was active, and those of the first and third DI muscles were 0.355 and 0.668, respectively, when the index and middle fingers were not active. The results implied that *f*_peak_ and *P*_max_ are not reliable enough to be used as the predictor variables of our logistic regression analysis among the 10 subjects in our experiment.

The mean SNRs of the frequency-domain variables (*f*_med_ and *f*_peak_) were lower than those of the other variables (e.g., 1.201 of *f*_med_ and 1.926 of *f*_peak_ of the first DI muscle). The results also indicated a subtle distinction between the active and non-active states.

Fig. 4 shows the box plots of the mean amplitudes (*V*_mean_) of all DI muscles when one of the fingers was active. The results showed that the distribution of the amplitude of the DI muscles inserted into the active fingers was higher than that of the non-active fingers. Table 3 also shows that the amplitude of the inserted muscles of the active fingers was significant, while that of the non-inserted muscles was not, even though some distributions of the amplitude seemed higher in Fig. 4 (e.g., the fourth DI muscle in Fig. 4 (d) when the little finger was active). This result implies the necessity of the validation of the significance. The results of the univariate logistic regression models whose predictor variables are the amplitude-domain (i.e. *V*_max_ in Table 2 and *V*_mean_ in Table 3) and the power-domain (i.e. *P*_max_ in Table 4 and *P*_total_ in Table 5) showed significant consistency between the activated and inserted fingers of the DI muscles into the proximal phalanx, as shown in Fig. 3.

**Figure 4.**
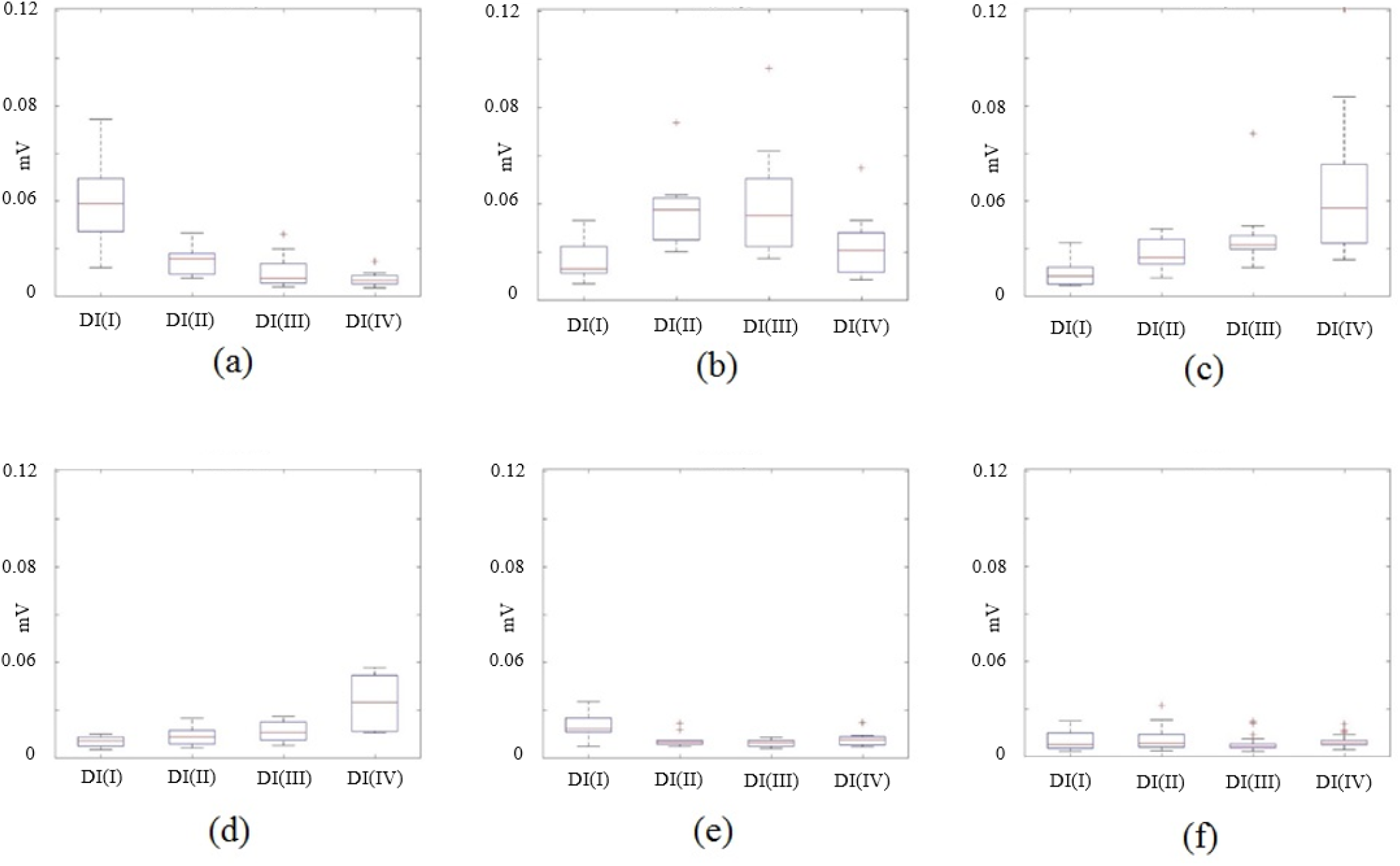
Box plots of the mean amplitudes (mV) of the sEMG signals of the first DI(I) to the fourth DI(IV). (a) Index as active finger. (b) Middle as active finger. (c) Ring as active finger. (d) Little as active finger. (e) Thumb as active finger. (f) No active fingers.

**Table 2.** *P*-values to predict the active and non-active states of each finger (the predictor variable is *V*_max_ in Table 1).

| Finger | $P$-value ($V_{\max}$) | | | |
| --- | --- | --- | --- | --- |
|  | DI(I) | DI(II) | DI(III) | DI(IV) |
| Index | <0.001 | 0.611 | 0.822 | 0.159 |
| Middle | 0.399 | <0.001 | <0.001 | 0.171 |
| Ring | 0.436 | 0.133 | 0.012 | <0.001 |
| Little | 0.194 | 0.458 | 0.776 | 0.088 |
| Thumb | 0.695 | 0.112 | 0.153 | 0.210 |

**Table 3.** *P*-values to predict the active and non-active states of each finger (the predictor variable is *V*_mean_ in Table 1).

| Finger | $P$-value ($V_{\text{mean}}$) | | | |
| --- | --- | --- | --- | --- |
|  | DI(I) | DI(II) | DI(III) | DI(IV) |
| Index | <0.001 | 0.344 | 0.749 | 0.209 |
| Middle | 0.292 | <0.001 | <0.001 | 0.128 |
| Ring | 0.557 | 0.113 | 0.012 | <0.001 |
| Little | 0.159 | 0.351 | 0.832 | 0.088 |
| Thumb | 0.680 | 0.173 | 0.203 | 0.232 |

**Table 4.** *P*-values to predict the active and non-active states of each finger (the predictor variable is *P*_max_ in Table 1).

| Finger | $P$-value ($P_{\max}$) | | | |
| --- | --- | --- | --- | --- |
|  | DI(I) | DI(II) | DI(III) | DI(IV) |
| Index | <0.001 | 0.323 | 0.704 | 0.190 |
| Middle | 0.395 | <0.001 | <0.001 | 0.169 |
| Ring | 0.476 | 0.123 | 0.016 | <0.001 |
| Little | 0.191 | 0.346 | 0.947 | 0.036 |
| Thumb | 0.406 | 0.189 | 0.204 | 0.241 |

**Table 5.** *P*-values to predict the active and non-active states of each finger (the predictor variable is *P*_total_ in Table 1).

| Finger | $P$-value ($P_{\text{total}}$) | | | |
| --- | --- | --- | --- | --- |
|  | DI(I) | DI(II) | DI(III) | DI(IV) |
| Index | <0.001 | 0.317 | 0.807 | 0.226 |
| Middle | 0.236 | <0.001 | <0.001 | 0.090 |
| Ring | 0.585 | 0.099 | 0.010 | <0.001 |
| Little | 0.183 | 0.377 | 0.841 | 0.119 |
| Thumb | 0.741 | 0.159 | 0.200 | 0.229 |

We also showed pseudo-*R*^2^ coefficients proposed for the logistic regression analysis (Nagelkerke, 1991). The Nagelkerke *R*^2^ coefficients used in our analysis are shown in Table 6. The range of Nagelkerke *R*^2^ coefficients is from 0.0 to 1.0, and intuitively easy to understand by imitating the range of the general non-pseudo ordinary least squares *R*^2^. The result represented a high correlation of DI muscles during the action of the inserted fingers, but a low correlation during the action of the non-inserted fingers. Meanwhile, the amplitudes of the signals of DI(I), which was inserted into the index finger, only indicated a high correlation during the action of the index finger (0.720). DI(II) and DI(III) were both inserted into the middle finger, and similarly, the results indicated a high correlation (0.779 and 0.508) during the action of the middle finger. However, the results of DI(IV) showed only a relatively high correlation, but lower than the formers (0.537), during the action of the ring finger. Also, none of the muscles displayed any high correlation during the action of the little finger and the thumb, which were not inserted into any DI muscle, even though the position of the fingers were adjacent to the DI muscles (i.e., DI(I) to the thumb and DI(IV) to the little finger).

**Table 6.** Nagelkerke *R*^2^ of the univariate analysis (the predictor variable is *V*_mean_ in Table 1).

| Finger | Nagelkerke $R^2$ ($V_{\text{mean}}$) | | | |
| --- | --- | --- | --- | --- |
|  | DI(I) | DI(II) | DI(III) | DI(IV) |
| Index | 0.720 | 0.017 | 0.002 | 0.068 |
| Middle | 0.020 | 0.779 | 0.508 | 0.042 |
| Ring | 0.009 | 0.047 | 0.133 | 0.537 |
| Little | 0.075 | 0.025 | 0.000 | 0.054 |
| Thumb | 0.003 | 0.063 | 0.066 | 0.057 |

As a comparison, we also showed a result of Student’s *t*-test in Table 7 as the *P*-values and Table 8 as the partial η^2^, in case that the predictor variable is *V*_mean_. We obtained similar results of the *P*-values between Table 3 and 7, and the decision coefficients between Table 6 and 8. However, the *P*-value of DI(III) in Table 7, which is adjacent and not inserted to the metacarpal bone of the ring finger, is quite low (0.001) when the ring finger is active, even though the *P*-value in Table 3 is much higher (0.012) than the threshold (0.001).

**Table 7.** *P*-values of Student’s *t*-test (the predictor variable is *V*_mean_ in Table 1).

| Finger | $P$-value ($V_{\text{mean}}$) | | | |
| --- | --- | --- | --- | --- |
|  | DI(I) | DI(II) | DI(III) | DI(IV) |
| Index | <0.001 | 0.336 | 0.751 | 0.207 |
| Middle | 0.280 | <0.001 | <0.001 | 0.094 |
| Ring | 0.555 | 0.088 | 0.001 | <0.001 |
| Little | 0.168 | 0.350 | 0.833 | 0.054 |
| Thumb | 0.682 | 0.172 | 0.206 | 0.232 |

**Table 8.**
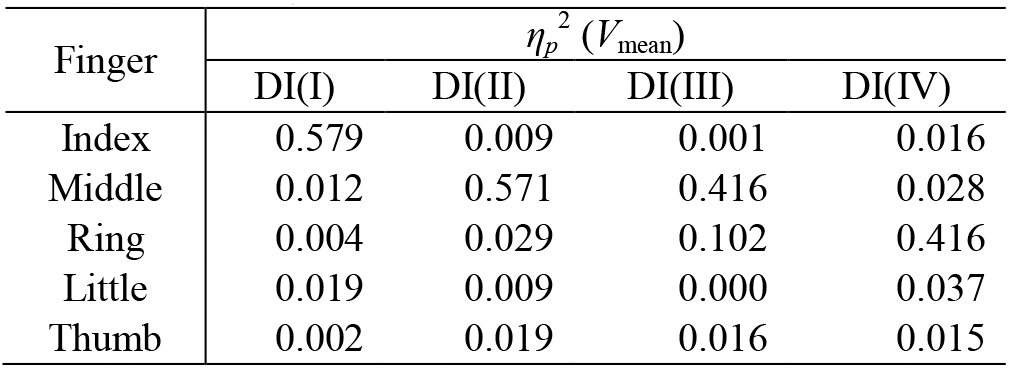
Partial *η*^2^ of Student’s *t*-test (the predictor variable is *V*_mean_ in Table 1).

| Finger | $\eta_p^2$ ($V_{\text{mean}}$) | | | |
| --- | --- | --- | --- | --- |
|  | DI(I) | DI(II) | DI(III) | DI(IV) |
| Index | 0.579 | 0.009 | 0.001 | 0.016 |
| Middle | 0.012 | 0.571 | 0.416 | 0.028 |
| Ring | 0.004 | 0.029 | 0.102 | 0.416 |
| Little | 0.019 | 0.009 | 0.000 | 0.037 |
| Thumb | 0.002 | 0.019 | 0.016 | 0.015 |

### 3.2. Multivariate Analysis

We analyzed a multivariate logistic regression model that consisted of two types of predictor variables in two ways: *V*_mean_- *V*_max_ and *P*_total_-*f*_med_. The selections were made by combining the integrated power (or amplitude) with the peak power (*V*_mean_-*V*_max_) and the power domain with the frequency domain (*P*_total_-*f*_med_). *f*_peak_ and *P*_max_ are not listed because the ICCs shown in Table 1 are relatively lower than other predictor variables, as we mentioned in Section 3.1. *V*_mean_ was also omitted from the combination with any frequency-domain variable because the non-normalized mean amplitude was equivalent to the integrated power (*P*_total_). The results are shown in Tables 9 as Nagelkerke *R*^2^. The correlation values of univariate regression models using *V*_mean_ and *P*_total_ are also included in the tables as a comparison to those of multivariate regression models.

**Table 9.** Nagelkerke *R*^2^ of the multivariate analysis.

| Finger<br>Electrode<br>Position | Variables |  |  |  |
| --- | --- | --- | --- | --- |
| | $V_{\text{mean}}$ | $V_{\text{mean}} - V_{\text{max}}$ | $P_{\text{total}}$ | $P_{\text{total}} - f_{\text{med}}$ |
| Index |  |  |  |  |
| DI(I) | 0.720 | 0.729 | 0.712 | 0.716 |
| Middle |  |  |  |  |
| DI(II) | 0.779 | 0.795 | 0.775 | 0.777 |
| DI(III) | 0.508 | 0.529 | 0.515 | 0.555 |
| DI(II,III) | 0.783 | 0.833 | 0.775 | 0.799 |
| Ring |  |  |  |  |
| DI(IV) | 0.537 | 0.574 | 0.583 | 0.716 |

The results of Nagelkerke *R*^2^ in Table 9 from the correlation of the multivariate logistic regression models (i.e., *V*_mean_-*V*_max_ and *P*_total_-*f*_med_ as the predictor variables) were slightly higher than the results of the univariate logistic regression models (i.e., *V*_mean_ and *P*_total_ as the predictor variables). The models with two muscles are listed in the tables as DI(II,III) during the action of the middle finger because of the two insertions of the DI muscles into the middle finger. However, the correlation was not necessarily higher than the correlation of DI(II). Only the multivariate regression models of the ring finger showed a higher increase (0.716 with *P*_total_-*f*_med_) as compared to the univariate regression models (0.537 with *V*_mean_ and 0.583 with *P*_total_).

The results of the sensitivity and the specificity of the univariate and multivariate regression models are shown in Table 10. These measures of the binary classification were computed with the mean values of the samples of the same experimental session of each subject, with the fact that the ICCs of all predictor variables used in Table 10 were high enough (at least 0.918 of *P*_total_ at the index-finger action in Table 1). It can be seen that only the sensitivity of the discrimination of the activity of the ring finger is apparently improved from 0.5 to 0.8 while the specificity is not decreased, when the multivariate model with *P*_total_-*f*_med_ was applied instead of the univariate model with *P*_total_.

**Table 10.**
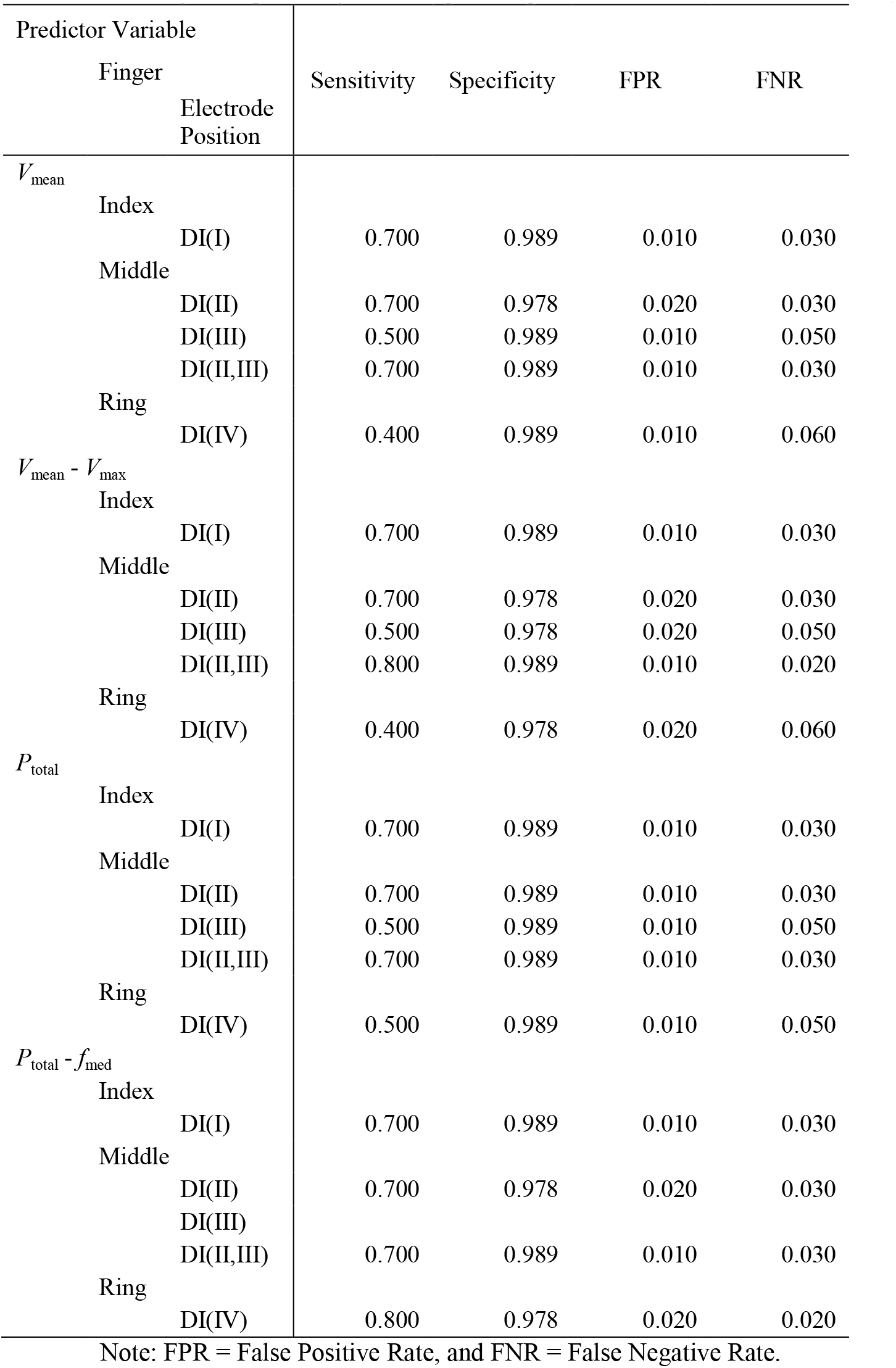
Sensitivity and specificity of the logistic regression models (cutoff-value = 0.5).

As a result, we hypothesized that the regression models of the forcing action of the ring finger had more complexity than the other two fingers with the insertions of the DI muscles (i.e., the index finger and the middle finger), which was also contributed by some muscle weaknesses and inexperienced actions. Our experimental results showed that the combination of the predictor variables *P*_total_ (or *V*_mean_ as the equivalent) and *f*_med_ in the frequency domain were the most important key predictor variables of our regression models. Further experiments with more participants might be needed to verify the hypothesis of the multivariate models because some of the ICCs of the predictor variables shown in Table 1 were reasonably low, as mentioned in the section of univariate analysis (i.e., 0.509 at DI(III) with *f*_peak_, 0.355 at DI(I), and 0.668 at DI(III) with *P*_max_).

## Conclusions

In this paper, we evaluated activities of densely located interosseous hand-muscles during finger-oriented tasks through analyzing the surface electromyographic signals using univariate and multivariate logistic regression models. Ten healthy subjects participated in our experiment. The results of the univariate analysis with the power and amplitude domain predictor variables of the surface electromyographic signals showed significant consistency between the activated and inserted fingers of the DI muscles into the proximal phalanx (*P* < 0.001). Meanwhile, the results of the multivariate analysis showed a higher correlation of the regression model of the fourth DI muscle during the action of the ring finger using frequency-domain variables (Nagelkerke *R*^2^ = 0.716 when the median frequency was used), compared to the model without the frequency-domain variables (Nagelkerke *R*^2^ = 0.583). Our result showed that the logistic regression models have a particular possibility for the analysis of densely located hand-muscles using the surface electromyography which is non-invasive and safe to use, and may promote researching variable finger activities statistically.

## Acknowledgement

The authors would like to thank Ryohei Koyama, Prof. N. Togawa, and Prof. Y. Shi for their contributions to this work.

